# Learning universal knowledge graph embedding for predicting biomedical pairwise interactions

**DOI:** 10.1101/2025.02.10.637419

**Authors:** Siyu Tao, Yang Yang, Xin Liu, Yimiao Feng, Jie Zheng

## Abstract

Predicting biomedical interactions is crucial for understanding various biological processes and drug discovery. Graph neural networks (GNNs) are promising in identifying novel interactions when extensive labeled data are available. However, labeling biomedical interactions is often time-consuming and labor-intensive, resulting in low-data scenarios. Furthermore, distribution shifts between training and test data in real-world applications pose a challenge to the generalizability of GNN models. Recent studies suggest that pre-training GNN models with self-supervised learning on unlabeled data can enhance their performance in predicting biomedical interactions. Here, we propose LukePi, a novel self-supervised pre-training framework that pre-trains GNN models on biomedical knowledge graphs (BKGs). LukePi is trained with two self-supervised tasks: topology-based node degree classification and semantics-based edge recovery. The former is to predict the degree of a node from its topological context and the latter is to infer both type and existence of a candidate edge by learning semantic information in the BKG. By integrating the two complementary tasks, LukePi effectively captures the rich information from the BKG, thereby enhancing the quality of node representations. We evaluate the performance of LukePi on two critical link prediction tasks: predicting synthetic lethality and drug-target interactions, using four benchmark datasets. In both distribution-shift and low-data scenarios, LukePi significantly outperforms 15 baseline models, demonstrating the power of the graph pre-training strategy when labeled data are sparse.

## I. Introduction

**N**UMEROUS biomolecules within living cells form diverse interactions, acting as pathways or direct participants in various biological activities and functions [4]. For example, protein-protein interactions (PPIs) are essential for many biological processes and cellular functions [27]. Discovering biological interactions is crucial for understanding cellular behaviors and vital for disease prevention, diagnosis, treatment, and drug discovery [6], [37]. Experimental screening is reliable but hindered by high costs and time-consuming procedures, etc. By contrast, machine learning approaches can significantly reduce the search space, thereby accelerating the discovery of new interactions [2], [20], [31]. Consequently, machine learning techniques have played increasingly important roles in predicting biological interactions. Moreover, real-world biomedical data are commonly represented in the form of graphs. In particular, biomedical knowledge graphs (BKGs) are well-suited for modeling complex interactions in biological systems, integrating various types of data source [6], [38]. In these graphs, nodes represent biomedical entities (e.g., proteins, RNAs, and drugs), while edges represent the interactions between nodes, such as drug-drug interactions. Consequently, predicting biological interactions is commonly formulated as the link prediction task, where the objective is to classify potential links between biomedical entities as either present or absent. Here, we focus on the prediction of two types of pairwise interactions: synthetic lethality (SL) and drug-target interactions (DTIs). Specifically, SL is a prominent type of genetic interaction that occurs when the inhibition of two genes is lethal, while the inhibition of each single gene is not [28], [31]. DTI refers to the interaction between a drug and its target protein, serving as an important step in the process of drug discovery [2], [32].

In biomedical networks, graph neural networks (GNNs), a popular deep learning model for graph-structured data, are widely used for biological link prediction tasks [2], [5], [8], [19], [20], [31], [36]. For example, KG4SL [31], the first model to predict SLs by learning from a BKG, uses a GNN that samples 2-hop neighbors of genes from the BKG to capture local graph structures around the genes. MVGCN-iSL [8] is a multi-view graph auto-encoder model that incorporates gene features derived from five networks, including an SL graph, a protein-protein interaction (PPI) network and a gene co-expression network. GNN-based approaches have shown potential in predicting DTIs. For instance, GraphDTA [22] is developed to predict DTIs by constructing drug-related graphs to represent drug molecules and applying multiple GNNs to extract molecular features. DrugBAN [2] is a deep bilinear attention network which employs GCNs to encode local structures of 2D molecular graphs, generating representations for both drugs and proteins. Although these GNN-based methods have exhibited promising potential in predicting biological pairwise interactions, some limitations persist. First, training a GNN model for a specific interaction prediction task is typically conducted in a supervised manner, necessitating sufficient label data to ensure an effective training process. However, obtaining sufficient task-specific labels is often infeasible due to the high costs and time-consuming process of labeling biomedical data [21]. As such, the scarcity of labeled data, known as low-data scenarios, often leads to suboptimal performance of the supervised models. Secondly, owing to the vast regions of pairwise biological molecular space, these models also need to predict interactions involving new nodes, which are often unseen in the training data. These interactions have different distributions from that of the training data, and therefore leads to a distribution-shift issue. However, supervised training of deep GNN-based models on these restricted datasets may result in poor generalizability of the models, hindering their application in real-world scenarios [2], [13].

To alleviate these predicaments, a widely-used strategy is self-supervised pre-training, which involves learning from abundant unlabeled data and then fine-tuning the pre-trained model on specific downstream tasks [32], [34]. This learning process does not require labeled data, making it advantageous for tasks with limited labeled data. For instance, Edge-Mask [13] randomly masks edges and trains a GNN model to predict the types of the masked edges. MSSL2drug [32], a recent graph-attention-based multitask adversarial learning framework, enhances DTI prediction by leveraging multitask self-supervised learning on BKGs and exploring 15 combinations of popular self-supervised tasks. However, the objectives of pre-training tasks and downstream tasks are often different, which may result in the inability to capture the specific features or knowledge required for predicting biological interactions during the pre-training phase [24]. Consequently, such models may underperform in certain interaction prediction tasks. Therefore, designing effective pre-training tasks to improve the performance of various biological link predictions, particularly in distribution-shift and low-data scenarios, remains a challenge.

Biological systems are highly interdependent, and the formation of specific interactions is often influenced by others. For example, SL is closely associated with PPIs [28]. Since BKGs integrate various biological interactions, we hypothesize that capturing the shared knowledge underlying these interactions can provide rich supplemental information for learning effective representations. This shared knowledge may reduce the model’s reliance on specific interaction information, improving its performance in low-data scenarios. By leveraging abundant biomedical knowledge for pre-training, models can enhance their ability to learn generalizable representations, thereby improving performance in distribution-shift scenarios. Furthermore, pre-training on BKGs offers the potential for models to be applied across diverse biological link predictions. Here, we propose LukePi (**L**earning **u**niversal **k**nowledge graph **e**mbedding for predicting biomedical **P**airwise **i**nteractions), a novel self-supervised pre-training framework designed to enhance biomedical interaction predictions. The key idea of LukePi is to pre-train a GNN model on BKGs by simultaneously employing two simple yet powerful self-supervised tasks. Specifically, we introduce two self-supervised pre-training tasks tailored to biomedical interactions on a BKG, namely topology-based node degree classification and semantics-based edge recovery. First, we compute node degrees in the original BKG and construct a disrupted BKG by masking edges and adding pseudo edges. The node degree classification task predicts node degree labels by using a GNN model on a disrupted BKG derived from the original BKG, increasing the difficulty of the task. This challenging task forces the GNN model to focus on and learn the local information of nodes, thereby extracting robust biological patterns contained in the topological structure. Secondly, in addition to topological information, abundant semantic information provided in BKGs also enables the GNN model to achieve a fine-grained understanding of various interactions. Consequently, the edge recovery task is developed to help the model capture semantic features and associations among various interactions in the disrupted BKG by predict both the type and existence of candidate edges. This dual-task pre-training strategy provides LukePi with a comprehensive view, enabling the model to effectively capture both topological and semantic information, thus improving the effectiveness and generalizability of node representations. To verify the effectiveness of LukePi, we compare it with 15 baselines on two downstream tasks, i.e. SL prediction and DTI prediction, across various evaluation scenarios. Results show that LukePi significantly outperforms these baselines, demonstrating its strong effectiveness and generalizability in distribution-shift and low-data scenarios. Additionally, case studies of the predicted SL pairs, including literature, biological mechanism analysis, and survival analysis, further demonstrate the potential of LukePi in real-world applications. Overall, the main contribution of this paper are summarized as follows:

- We propose LukePi, a novel self-supervised pre-training framework on biomedical knowledge graphs (BKGs) for predicting various biomedical pairwise interactions.
- LukePi introduces two powerful self-supervised tasks to capture both the topological and semantic characteristics of BKGs. This dual-task pre-training strategy enables effective node representations, thereby improving the accuracy of downstream interaction predictions.
- We systematically evaluate LukePi on two crucial link prediction tasks (synthetic lethality and drug-target interactions) across four benchmark datasets. Compared to fifteen baseline models, LukePi achieves the best overall performance, demonstrating its effectiveness and strong generalizability.
- Furthermore, case studies on the predicted SL pairs show that LukePi not only identifies known SL pairs but also predicts promising ones, highlighting its potential for real-world applications.

## II. Methodology

### A. Construction of pre-training graph

BKGs are denoted as 𝒢 = (𝒱, ℰ), where 𝒱 and ℰ are defined as the sets of nodes and edges, respectively. We construct a graph 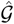 by modifying edges in the original BKG through a two-step process. Hereafter, we call 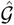 as the pre-training graph. This construction process consists of two transformation operations: topological transformation and semantic transformation. Firstly, we use the topological structure transformation to randomly sample pseudo edges as certain samples(i.e., those edges that do not exist in 𝒢). Note that the pseudo edges are also not added into the graph 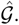 Meanwhile, we randomly drop some true edges in 𝒢 as another type of topological structure transformation. Then, we use the [MASK] symbol to replace the semantic features of the above true edges in 𝒢, which is called the semantic transformation. Figure 1 (**Step I**) illustrates the process of construction.

**Fig. 1.**
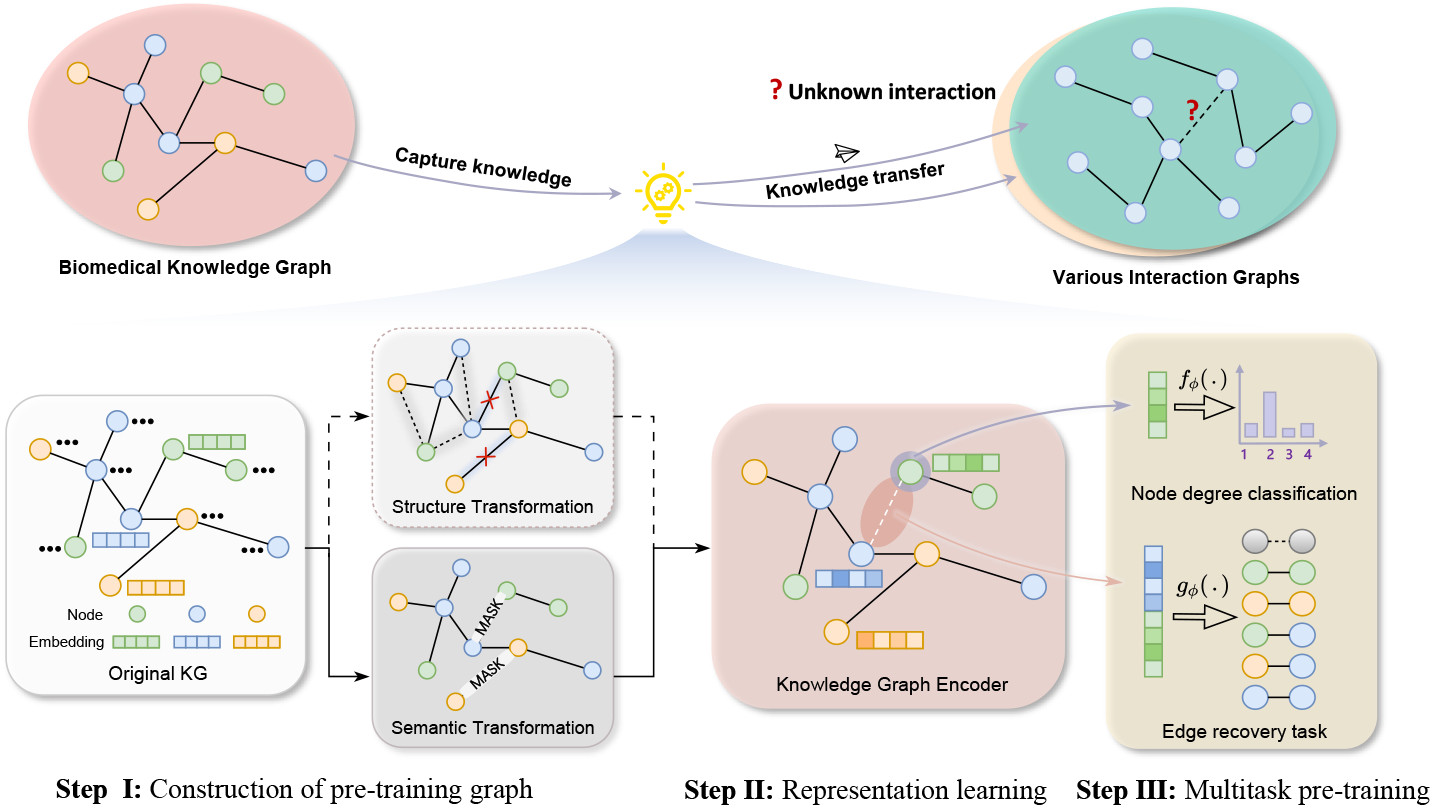
Overview of LukePi. LukePi is a pre-training framework based on two self-supervised tasks. **Step I**: Given a BKG, we calculate node degrees and classify them to generate degree labels. We then construct a pre-training graph 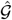 by modifying edges in the original BKG, identifying candidate edges for the edge recovery task. The topological transformation adds pseudo edges (gray dashed edges) and removes masked edges (red ‘×’). The semantic transformation masks the semantic feature of some real edges with the symbol [MASK]. **Step II**: LukePi uses a GNN model to embed the nodes of 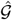, concatenating the pairwise node embeddings as candidate edge representations. **Step III**: These node and edge representations are input into two classifiers for task-specific predictions to train LukePi. Finally, LukePi is optimized by minimizing the combined loss from the two self-supervised tasks. After pre-training, LukePi captures shared knowledge to produce effective embeddings transferable to downstream interaction prediction tasks.

### B. Topology-based pre-training task of node degree classification

The importance of nodes in a biological network could reveal underlying mechanisms of biomolecular interactions [7]. As a basic topological attribute of nodes, the degree (i.e., the number of incident edges of a node) can indicate the potential importance of nodes. For example, protein TP53, a crucial gene in many cancers, is a high-degree node in PPI networks [1]. Considering the computational efficiency in handling large-scale graphs, we choose the degree value of each node as the objective because it is easy to compute. Therefore, we introduce a node degree classification task to guide the GNN model to learn from topological structures, enabling it to extract robust patterns that could enhance its generalizability.

The degree classification problem is formulated as a multiclass classification task. Specifically, given a BKG 𝒢, we first compute the degrees for all nodes and split them into 𝒬 intervals, i.e., a set of degree labels 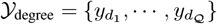, where larger label values correspond to nodes with higher degrees. To increase the challenge of this task, we apply the GNN model on the disrupted graph 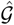, rather than the original BKG. The GNN encoder must fully learn the local neighboring topological information to accurately predict the degree label of each node. This comprehensive exploration of robust topological information could enhance the generation of effective node representations. Finally, a multi-layer perceptron (MLP) classifier takes the node representations **z** as input and predicts the degree label. The process is summarized as follows:

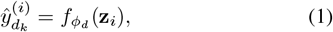

where 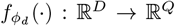 is the classifier, and **z**_*i*_ ∈ ℝ^*D*^ is the node embedding of each node v_*i*_. The output 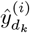 is a class probability vector, where each dimension corresponds to a class of node degrees, and the value indicates the probability of a node v_*i*_ belonging to that class.

### C. Semantics-based pre-training task of edge recovery

While the node degree classification task enables GNN models to extract robust topological information, relying solely on this information cannot distinguish the different biological information underlying similar topological graphs. To overcome this limitation, incorporating semantic knowledge along with topological information from BKGs allows GNN models to achieve a more fine-grained understanding of biological entities and processes, thereby significantly enhancing the accuracy of predicting biological interactions. To this end, we introduce a novel edge recovery task to learn how different types of interactions relate to each other within the BKG.

Specifically, after constructing the pre-training BKG, a candidate edge set is consisted by two parts of edges, i.e., 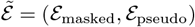. The introduction of pseudo edges enables the GNN model to distinguish between true and pseudo edges, thereby capturing the inherent features of the real edges. Moreover, the pseudo edges enrich the training dataset with diverse samples, pushing the model to gain a more comprehensive understanding of the semantic features of different interactions. Each candidate edge 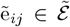 is represented by **h**_*ij*_ = [**z**_*i*_; **z**_*j*_], where **h**_*ij*_ ∈ ℝ^2*D*^ and [·; ·] is the concatenation operation. Notably, we add a “no-type” category to represent the pseudo edges, i.e., 𝒴_edge_ = {*y*_*s*1_, …, *y*_*sC*_} is expanded to 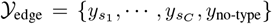 as labels for pre-training, where *R* is the number of edge classes. Previous edge-level masking pre-training strategies typically only predict the semantic features of the masked edges (i.e., the true edges of the graphs) [13], [32]. In contrast, our edge recovery task is more challenging, as it requires both the classification of the semantic features and the prediction of the existence of the edges. This dual requirement creates a more complex task than other self-supervised masking pre-training tasks, forcing the GNN models to better capture the regularities of the node and edge attributes distributed over the disrupted BKG 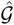. Finally, these edge representations are fed into a MLP classifier to perform multi-class classification. This process is expressed as:

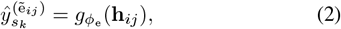

where 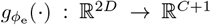 is the classifier. The output 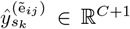 is a class probability vector, of which each element is the probability of the edge e^*ij*^ belonging to a distinct class.

### D. Knowledge graph encoder

Inspired by the advances of GNNs in graph representation learning [2], [31], we introduce a knowledge graph encoder to learn the node embeddings of 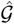. In the LukePi, we adopt the heterogeneous graph transformer (HGT) as the BKG encoder ℬ_*θ*_ [14], which is designed for heterogeneous graphs. Specifically, we first use Xavier’s uniform distribution to initialize the node representations 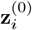 according to the category of nodes, i.e., nodes belonging to the same category have the same initial representation. Secondly, the node representation is processed in each HGT convolutional layer by three operators: heterogeneous mutual attention, heterogeneous message passing, and target-specific aggregation. By stacking *M* layers, followed by a type-specific linear projection **Linear**_*τ*_ and a residual connection [12], we obtain the node representations of all nodes in the graph 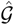. Finally, the representation of v_*i*_ is defined as follows:

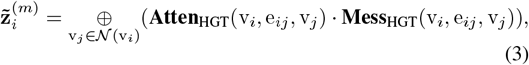

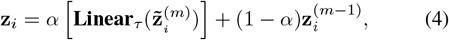

where 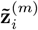 represents the embedding of the node v_*i*_ in the *m*-th layer. 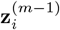 is the original vector of the previous *m*−1 layer. **Atten(**·) estimates the importance of each neighbor node, and **Mess(**·) extracts the messages from its neighbor node v_*j*_. ⊕ (·) averages the messages from the neighbor nodes *N* (v_*i*_) by the attention weight. *α* is a learnable parameter that adaptively determines the relative importance of the two components.

### E. Multitask joint pre-training

Previous studies have demonstrated that relying on a single self-supervised pre-training task may introduce bias to downstream interaction prediction tasks, harming the GNN model’s generalizability [32]. In contrast, pre-training on multiple self-supervised tasks could provide a more comprehensive view for the GNN model, enabling it to extract shared knowledge from a graph. Inspired by such observations, LukePi follows a multitask pre-training strategy.

Specifically, the training objective of LukePi consists of two components: the node degree classification and the edge recovery. We use a multi-class cross-entropy loss for both tasks and each task has a distinct classifier: 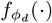 and 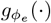, as shown in Eq.(1) and Eq.(2). Hence, the final pre-training loss is the sum of the losses from the two self-supervised pretraining tasks:

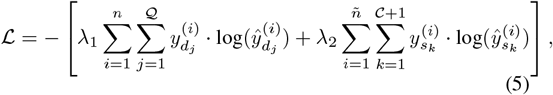

where the first term is the loss of the node degree classification task, and the second term represents the loss of the edge recovery task. Besides, *n* and ñ are the number of training samples in a batch and *λ*_1_ and *λ*_2_ are the coefficients for the losses of the two tasks.

## III. Experimental settings

### A. Datasets

#### 1) Pre-training dataset

Our pre-training dataset is the precision medicine knowledge graph (PrimeKG) [6], a comprehensive heterogeneous graph dataset covering a wide range of biomedical entities, which consists of 129,075 nodes and 8,100,498 edges. PrimeKG includes 10 types of nodes (e.g., protein, drug and pathway) and 30 types of edges (e.g., PPI and DTI). Moreover, we delete the interactions overlapping with the downstream datasets from the PrimeKG to prevent data leakage. More statistical details of PrimeKG are provided in Supplementary Tables S1 and S2.

### 2) Downstream task-specific datasets

For the SL prediction, we use two public datasets, i.e. SynLethDB [10] and an acute myeloid leukemia (LAML) SL dataset [11]. We use BindingDB [15] and Wang’s datasets [32] for DTI prediction. Specifically, SynLethDB is the most widely-used dataset of SL prediction, comprising 34,023 SL pairs and 2,900 non-SL pairs involving 9,845 genes. The LAML SL dataset is collected through the experimental screens. Following previous studies [8], [21], [29], [31], we randomly selected gene pairs that have not been confirmed as SL as negative samples so that the numbers of positive and negative labels are equal. BindingDB is a database of experimentally validated binding affinities, focusing primarily on DTI prediction. Followed by [2], we assign a label of 1 when the binding affinity score is below 100, and a label of 0 when the binding affinity score exceeds 10000. As such, we get an imbalanced dataset with 6,069 DTI pairs, including 4,527 negative pairs and 1,541 positive pairs. The Wang’s datasets [32] contains 4,978 positive samples and 4,994 negative DTI pairs. Table I provides the statistics of the four downstream datasets

**TABLE I.** Statistics of the three downstream datasets.

| Datasets | Node | Positive | Negative |
| --- | --- | --- | --- |
| SynLethDB [10] | 9,845 | 34,023 | 2,900 |
| LAML-specific SL dataset [11] | 261 | 995 | 995 |
| Drug target interaction [15] | 2,337 | 1,541 | 4,527 |
| Wang’s DTI dataset [32] | 4,978 | 4,994 | 2,524 |

### B. Evaluation scenarios

To evaluate LukePi comprehensively, we employ three data splitting strategies:

- **Vanilla splitting**: We randomly split the gene pairs into the training and test data.
- **Weak cold start**: One node of a gene pair from the test data is not present in the training data.
- **Cold start**: Neither of the nodes of a gene pair from the test data appears in the training data.

Besides, we adopt a split ratio of 8:2 for the training and test data in all data splitting settings. We use 5-fold crossvalidation (CV) for each scenario.

### C. Baselines

We compare LukePi with 15 representative GNN-based baselines for pairwise interaction predictions. These competitive baselines can be classified into three categories. First, we evaluate five GNN models: GCN [18], GAT [30], GIN [36], RGCN [26], and HGT [14]. Secondly, we introduce three self-supervised pre-training methods: EdgePred [13], Edge-Mask [13], and GCC [23]. Third, we evaluate seven task-specific methods: KG4SL [31] and PT-GNN [21] for the SL task, ELISL [29] and MVGCN-iSL [8] for the LAML-specific SL task; and GraphDTA [22], DrugBAN [2], and MSSL2drug [32] for the DTI task. In addition, the first eight methods [13], [14], [18], [23], [26], [30], [36] use PrimeKG as input data, while the seven task-specific methods use their original input data [2], [8], [21], [29], [31], [32]. It is worth noting that, since we treat SynLethDB data as pan-cancer data in this study, ELISL [29] and MVGCN-iSL [8] designed to predict potential SLs in specific cancers, are not included in the following comparisons using SynLethDB. We employ four well-known metrics to quantitatively evaluate the model performance. These metrics include area under the receiver operating characteristic curve (AUC), area under the precision-recall curve (AUPR), F1 score and balanced accuracy (BACC). Note that their values range from [0, 1], where higher values indicate better performance. The code of the baselines are obtained from the GitHub open source, and the detailed introduction of baselines are provided in Supplementary Material.

### D. Implementation details

#### 1) Pre-training configurations

LukePi adopts an HGT model [14] as the GNN encoder. The model is implemented using the Pytorch framework. All the experiments were conducted on one Nvidia V100 GPU. Specifically, we set the dimension of node representations to 128. The HGT encoder consists of four layers of HGT blocks, where the sizes of the layers are [256, 128], and the number of attention heads is set to 4. During pre-training, LukePi was trained using a batch size of 128 for 15 epochs with the Adam optimizer [17]. Since the pre-training dataset is of Web-scale, we adopt HGSampling [14] with four sampled layers to obtain sub-graphs, sampling 1024 nodes for each layer. For the edge recovery task, we set the hyperparameter mask rate *α* to 0.2 and used a 2-layer MLP classifier *g*_*ϕ*_(·) with a size of 128. For the node degree task, we set the hyperparameter degree label 𝒬 to 9, using a 2-layer MLP classifier *f*_*θ*_(·) with a size of 128. Both MLPs use ReLU as the activation function. To optimize the two MLP classifiers, we employed two Adam optimizers. The coefficients *λ*_1_ and *λ*_2_ were both set to 1. The learning rates for all pre-training stages were set to 1e-3.

#### 2) Fine-tuning configurations

When the fine-tuning of LukePi on downstream tasks, we used a 4-layer MLP classifier with sizes of [128, 64, 32]. After each linear layer, we applied a ReLU activation function. The batch size was set to 512, and the training was conducted for 100 epochs for the SL task in the vanilla splitting scenario. In other scenarios, the training was limited to 50 epochs. To prevent overfitting, a dropout rate of *β* = 0.2 was introduced. In the vanilla splitting scenario, the HGT encoder and the MLP classifier were optimized using two separate Adam optimizers with learning rates of 1e-1 and 1e-3, respectively. For weak cold start and cold start scenarios, the HGT encoder was frozen, and only the 4-layer MLP classifier was fine-tuned with a learning rate of 1e-3

## IV. Results

### A. Performance in the vanilla splitting scenarios

We first briefly demonstrate the performance of LukePi in the vanilla splitting scenarios. Figure 2A shows the average AUC scores of LukePi and other methods in these scenarios. LukePi exhibits overall superior performance in the SL and DTI tasks, though it only slightly lags behind the DTI task-specific methods. This is not unexpected, as task-specific models can fully leverage their potential when sufficient data are available. LukePi aims to improve performance in biological link predictions in distribution-shift and low-data scenarios, which are common in real-world applications. Although LukePi does not achieve the absolute best performance in both link prediction tasks, it remains competitive with task-specific methods in the vanilla scenarios.

**Fig. 2.**
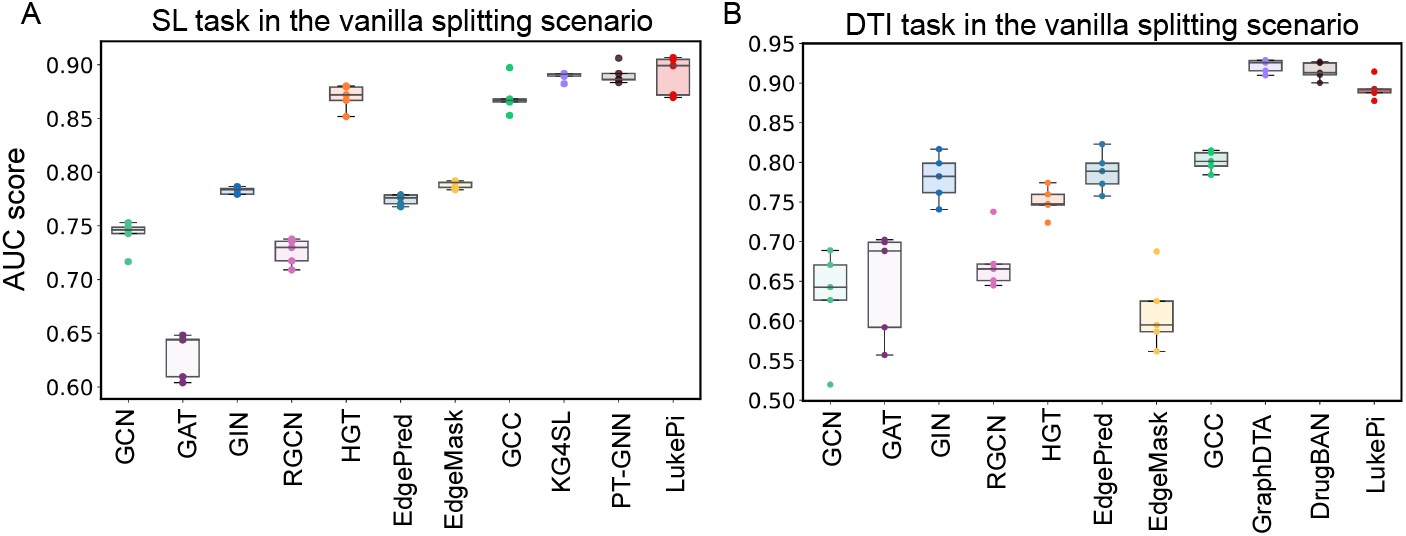
Comparison of LukePi with baselines in the vanilla splitting scenario. (A) SL prediction results. (B) DTI prediction task results.

### B. Improvement in distribution-shift scenarios

The promising performance of LukePi in the vanilla splitting scenario motivates us to further evaluate it in challenging distribution-shift scenarios. Consequently, we conduct experiments on two distribution-shift scenarios of varying difficulty: weak cold start and cold start. As shown in Table II and Table III, there are two main observations. First, all the models suffer from performance degradation in the two distribution-shift scenarios, compared to the vanilla data splitting. Notably, the four task-specific methods yield moderate results despite their sophisticated design for predicting specific interactions. These results clearly demonstrate the challenge posed by the distribution shift, where the nodes in the test data are unseen in the training data. Secondly, LukePi achieves the best performance in all scenarios. For example, LukePi outperforms the second-best methods with AUC improvements of +2.91% (SL) and +11.73% (DTI) in the cold start scenarios, respectively. The impressive performance of LukePi in distribution-shift scenarios could be attributed to the integration of effective self-supervised tasks. The two key tasks provide a comprehensive perspective for LukePi to capture shared knowledge across various interactions, thereby enabling the generation of powerful embeddings.

**TABLE II.** Comparison of LukePi with SL prediction methods in the weak cold start and cold start scenarios. The results (mean±std.) are reported based on 5-fold CV. The best performance and the runner-up performance are highlighted in bold and underlined, respectively.

| Method | Weak Cold Start (SL) |  |  |  | Cold Start (SL) |  |  |  |
| --- | --- | --- | --- | --- | --- | --- | --- | --- |
|  | AUC (%) | AUPR (%) | F1 (%) | BACC (%) | AUC (%) | AUPR (%) | F1 (%) | BACC (%) |
| GCN [18] | 70.33±2.38 | 69.92±2.47 | **68.13±0.47** | 56.93±1.83 | 65.72±6.28 | 66.16±5.27 | **65.75±4.79** | 57.10±3.33 |
| GAT [30] | 60.07±2.52 | 59.68±2.56 | 59.12±8.26 | 53.91±3.34 | 58.63±3.74 | 59.15±4.62 | 64.01±5.15 | 50.54±1.22 |
| GIN [36] | 70.55±2.02 | 73.91±1.47 | 60.46±5.24 | 65.06±1.70 | 67.55±5.42 | 71.44±5.47 | 48.70±12.25 | 61.69±5.23 |
| RGCN [26] | 68.10±1.69 | 67.54±1.07 | 63.83±4.18 | 64.73±1.81 | 60.18±4.40 | <u>59.21±2.77</u> | 58.11±7.55 | 60.68±5.20 |
| HGT [14] | 75.31±1.98 | 77.04±1.46 | 65.85±3.12 | 69.32±1.79 | 69.19±0.07 | 69.75±0.04 | 56.62±0.10 | 60.44±0.05 |
| KG4SL [31] | 69.75±2.85 | 67.57±1.26 | 54.63±5.63 | 63.42±2.44 | 55.22±5.33 | 54.78±4.13 | 33.08±4.72 | 52.81±2.65 |
| PT-GNN [21] | 63.02±2.56 | 67.26±3.57 | 38.37±6.85 | 59.80±2.86 | 49.55±5.82 | 50.23±3.06 | 23.00±0.31 | 50.00±0.12 |
| EdgePred [13] | 71.81±1.80 | 73.62±1.32 | 63.68±3.47 | 65.12±1.45 | 68.38±5.31 | 71.23±3.17 | 61.16±6.18 | 61.67±3.54 |
| EdgeMask [13] | 73.92±1.98 | 74.83±1.43 | 63.89±2.72 | 66.97±1.80 | 69.29±6.89 | 71.43±5.18 | 58.74±7.98 | 63.30±5.59 |
| GCC [23] | <u>76.46±0.02</u> | <u>77.56±0.01</u> | 67.63±0.04 | 70.17±1.06 | <u>67.72±0.48</u> | 71.05±0.25 | 58.27±0.58 | <u>62.86±0.33</u> |
| LukePi(Ours) | **78.73±2.18** | **78.88±1.04** | <u>67.94±0.13</u> | **71.21±0.06** | **72.20±7.55** | **74.36±4.85** | 65.13±6.52 | **65.99±6.27** |

**TABLE III.** Comparison of LukePi with DTI prediction methods in the weak cold start and cold start scenarios. The results (mean±std.) are reported based on 5-fold CV.

| Method | Weak Cold Start (DTI) |  |  |  | Cold Start (DTI) |  |  |  |
| --- | --- | --- | --- | --- | --- | --- | --- | --- |
|  | AUC (%) | AUPR (%) | F1 (%) | BACC (%) | AUC (%) | AUPR (%) | F1 (%) | BACC (%) |
| GCN [18] | 53.20±2.64 | 52.46±1.60 | 63.99±2.90 | 50.55±1.27 | 53.09±10.28 | 57.34±8.29 | **65.13±7.89** | 55.83±5.30 |
| GAT [30] | 50.24±3.38 | 52.44±4.68 | 27.32±32.15 | 50.16±0.33 | 57.06±10.79 | 64.09±8.71 | 36.00±30.12 | 51.11±2.22 |
| GIN [36] | 54.67±2.62 | 62.08±4.90 | 44.28±6.40 | 53.89±4.23 | 51.94±10.47 | 57.40±11.50 | 33.56±17.02 | 52.28±8.41 |
| RGCN [26] | 49.33±2.87 | 50.10±4.11 | 44.56±15.17 | 50.89±3.33 | 53.64±11.25 | 53.91±9.26 | 58.39±12.50 | 51.26±4.06 |
| HGT [14] | 54.45±3.48 | 65.77±3.93 | 54.90±6.43 | 51.80±2.44 | 59.21±9.47 | <u>70.62±3.50</u> | 61.88±4.94 | 55.89±5.45 |
| GraphDTA [22] | 61.96±4.35 | 59.24±4.61 | 58.04±5.90 | 58.78±4.27 | 52.60±12.20 | <u>55.80±11.25</u> | 39.70±14.97 | 55.57±5.26 |
| DrugBAN [2] | 60.76±3.57 | 58.68±4.04 | 56.67±5.48 | 58.06±3.76 | 66.73±10.18 | 69.02±13.29 | 47.43±14.42 | 61.41±7.29 |
| EdgePred [13] | 54.69±2.10 | 63.02±3.79 | 46.95±8.06 | 54.16±3.27 | <u>56.20±9.38</u> | 64.51±7.84 | 44.01±10.02 | <u>55.90±8.24</u> |
| EdgeMask [13] | 56.57±2.19 | 64.18±5.33 | 48.84±11.54 | 54.92±3.63 | 58.50±6.81 | 67.75±5.14 | 48.68±8.35 | 59.18±5.11 |
| GCC [23] | 58.32±2.54 | 67.01±1.98 | 56.05±3.88 | 55.72±1.57 | 54.18±10.61 | 62.70±5.57 | 48.23±9.16 | 52.22±7.79 |
| LukePi(Ours) | **74.58±3.27** | **77.62±1.99** | **71.63±0.24** | **67.72±3.77** | **78.46±3.22** | **78.22±6.60** | 64.70±6.52 | **68.42±6.52** |

Furthermore, we compare LukePi with a recent multi-task self-supervised approach called MSSL2drug [32], which also aims to address the distribution-shift issue in predicting DTIs. Due to the lack of specific computational details, we are unable to transfer MSSL2drug to PrimeKG. Thus, we conduct experiments on the BKG provided by MSSL2drug, which includes 3,046 biomedical entities and 111,776 edges. As depicted in Figure 3A, LukePi significantly outperforms MSSL2drug in both vanilla splitting and cold start scenarios. Furthermore, LukePi demonstrates superior computational efficiency. Specifically, using an Nvidia Tesla V100 GPU, LukePi completes the pre-training and fine-tuning processes in approximately 3 minutes and 20 seconds in the cold start scenario, which is 24 times faster than MSSL2drug (i.e., about 1 hour and 20 minutes), as shown in Figure 3B.

**Fig. 3.**
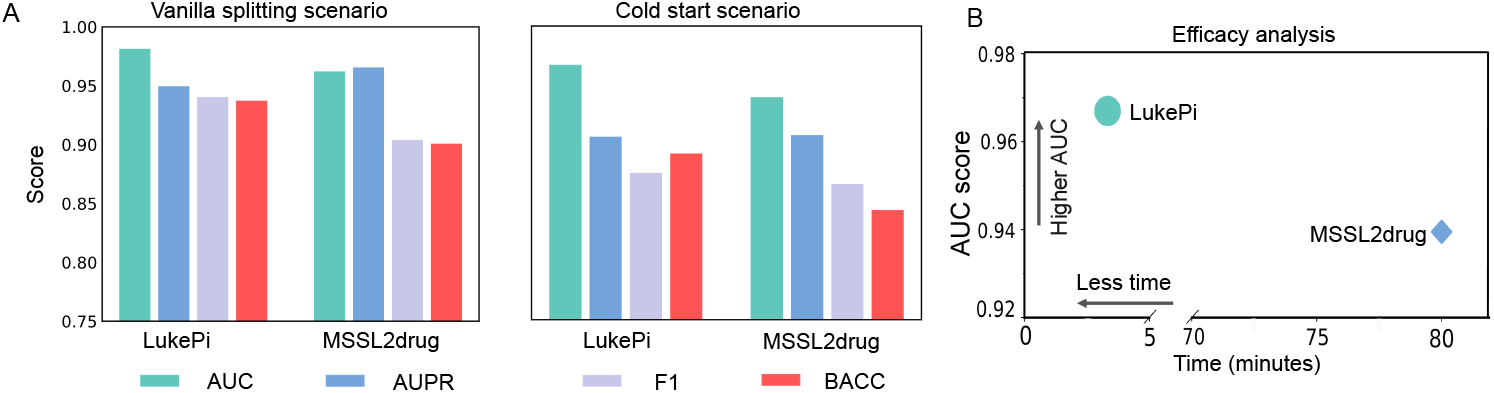
Comparison of LukePi with MSSL2drug [32]. (A) Left: the vanilla splitting scenario. Right: the cold start scenario. (B) Efficacy analysis. Comparison of the full required time (pre-training and fine-tuning) for LukePi and MSSL2drug, along with their test performance in the cold start scenario. The horizontal axis represents the required time and the vertical axis represents the average AUC score.

### C. Enhancement in the low-data scenarios

We proceed to evaluate the performance of LukePi under the low-data conditions. To simplify the experiments, we compare LukePi with seven representative methods, including GCN, HGT, EdgePred, EdgeMask, GCC, and all task-specific models (KG4SL and PT-GNN for the SL task; GraphDTA and DrugBAN for the DTI task). Then, we train the models with varying proportions of training data, ranging from 10% to 100%, for both SL and DTI tasks in the challenging weak cold start and cold start scenarios.

Figure 4 and Figure 5 illustrate the AUC scores for the SL and DTI tasks, respectively. We can make two observations: 1) The performance of all models degrades in the low-data scenarios, especially of the supervised methods (i.e. the GNN methods and the task-specific methods). For example, the performance of DrugBAN decreases significantly by 16.64% with 10% of the training data, compared to its performance using the full training data. 2) LukePi achieves the best overall performance across various training data ratios. For example, for the DTI task, LukePi can achieve better performance with only about 10% of the training data compared to all other methods using 100% of the training data, excepted for DrugBAN. We attribute the superior performance of LukePi in the low-data scenarios to its ability to incorporate shared interaction-related knowledge from PrimeKG. This valuable enhancement enables LukePi to generate effective embeddings, which is crucial in scenarios with limited label data. Moreover, the significant improvement of LukePi on the DTI prediction task, compared to the SL task, may be attributed to the fact that the pre-training BKG (PrimeKG) contains more drug-related information than genetic information.

**Fig. 4.**
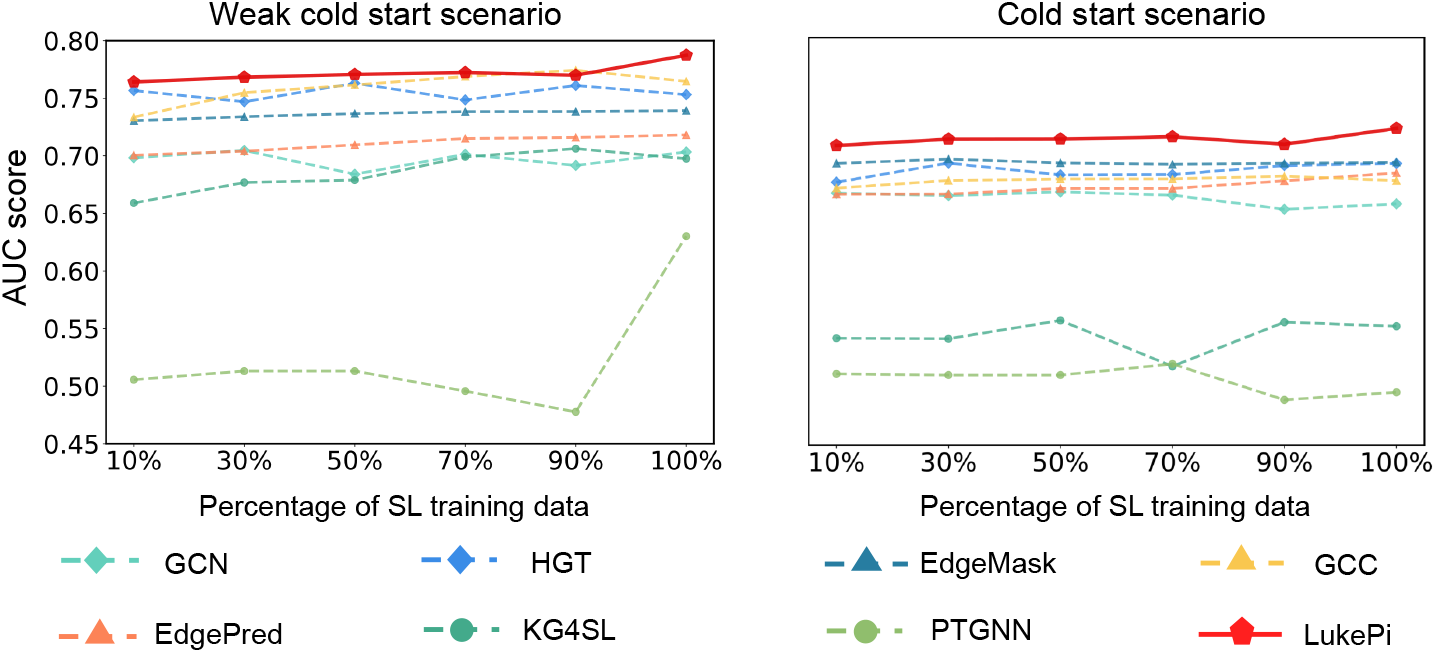
Comparison of LukePi with baseline models in the low-data scenarios (SL task). The horizontal axis represents the percentages of SL data for training, and the vertical axis represents the average AUC score.

**Fig. 5.**
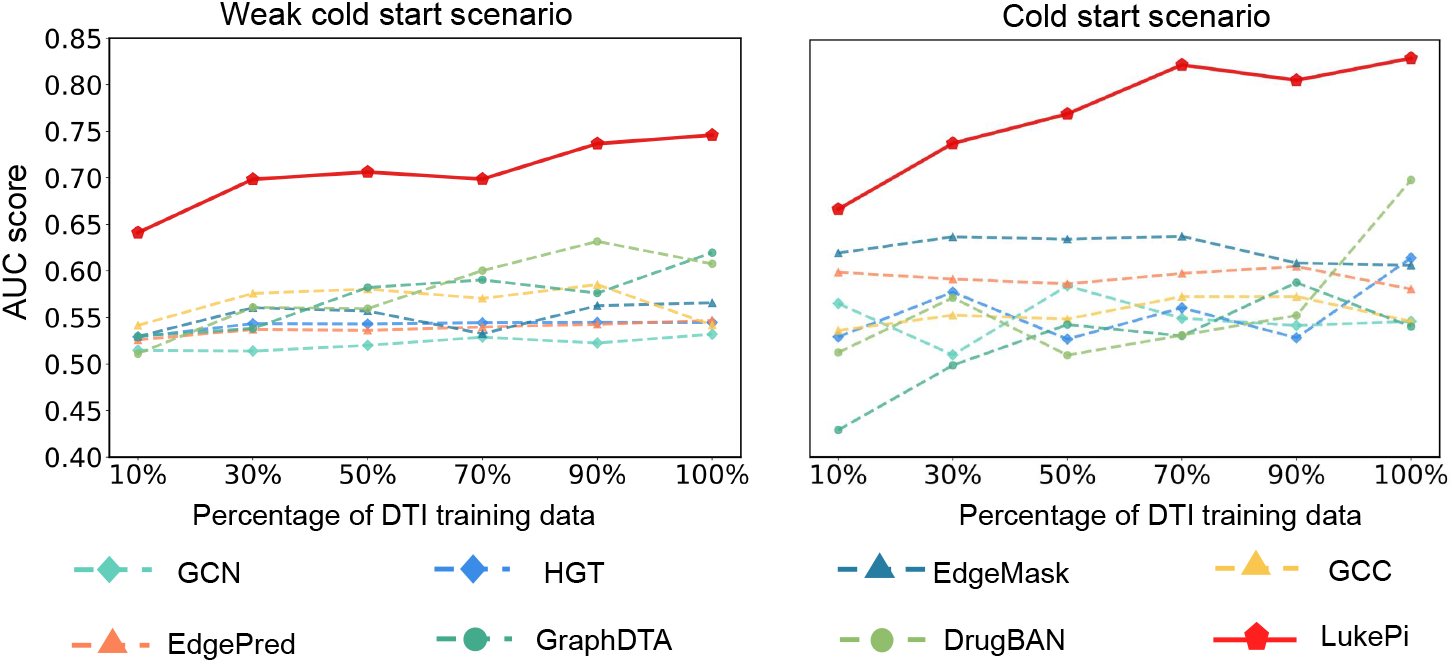
Comparison of LukePi with baseline models in the low-data scenarios (DTI task). The horizontal axis represents the percentages of DTI data for training, and the vertical axis represents the average AUC score.

Furthermore, as SLs have been found to be closely associated with genetic backgrounds [25], the prediction of SL interactions can be further refined for specific cancers. Notably, when focusing on a particular cancer type, the SL labels available tends to be sparse [25], which is considered a low-data scenario. To evaluate the performance of LukePi in this low-data scenario, we compare it with two recent context-specific SL prediction methods, that is, MVGCN-iSL [8] and ELISL [29] using an acute myeloid leukemia (LAML) SL dataset generated through experimental screening [11]. In addition, we compare LukePi with KG4SL and PTGNN, because they are aforementioned pan-cancer baseline models designed specifically for SL predictions. The dataset is divided into two subsets with a ratio of 8:2 for training and testing, respectively. The models are evaluated using 5-fold cross-validation, in the scenario of vanilla splitting. For this task of cancer-specific SL prediction, we do not evaluate the models in the scenario of distribution-shift, due to severe data sparsity which could lead to potential bias in the testing. We follow the original implementation details of LukePi, except for decreasing the learning rate from 1e-3 to 1e-4 and increasing the number of epochs from 30 to 40, in order to mitigate the overfitting. As shown in Table IV, LukePi achieves state-of-the-art performance in the LAML-specific SL prediction task. For example, LukePi achieves an improvement of +1.17% in the AUC score and +4.96% in the AUPR score compared with the second-best models. In conclusion, these results further demonstrate the superior performance of LukePi in the low-data SL prediction scenarios.

**TABLE IV.** Comparison of LukePi with SL prediction methods on a LAML dataset [11].

| Method | AUC (%) | AUPR (%) |
| --- | --- | --- |
| KG4SL [31] | 66.70±2.72 | 66.14±2.79 |
| PTGNN [21] | 64.86±1.97 | 65.41±2.26 |
| MVGCN-iSL [8] | 65.96±1.31 | 68.53±1.08 |
| ELISL [29] | 59.60±1.91 | 61.45±1.53 |
| LukePi(Ours) | **67.87±2.50** | **73.50±1.90** |

### D. Ablation study

To evaluate the effect of the two self-supervised tasks proposed in LukePi, we conduct an ablation study by creating three model variants: LukePi (without node degree classification task), which skips the node degree classification task; LukePi (without edge recovery task), which drops the edge recovery task, and LukePi (without pre-training), which is trained without any pre-training tasks. For both SL and DTI prediction tasks, we assess the performance of these three variants along with the original LukePi under weak cold start and cold start scenarios. We use the same experimental configurations to ensure a fair comparison. The results of the SL task and DTI task are presented in Table V and Table VI, respectively. The performance of two variants drops significantly compared with the full LukePi in each scenario. For example, in the SL prediction task, compared to the full LukePi, the average AUPR scores of LukePi (with the node degree classification task) and LukePi (with the edge recovery task) decrease by at least 3.52% and 0.86%, respectively. This decline is even more significant in the DTI prediction task. These results confirm the individual importance of the two pre-training tasks for the performance of LukePi. Secondly, integrating multiple pre-training tasks allows the model to learn more comprehensive representations, thereby enhancing its ability to adapt to different data distributions and scenarios. As shown in Table V and Table VI, only leveraging single pre-training tasks outperform LukePi (without pre-training) in some cases, but the improvement is inconsistent in specific scenarios. For example, for SL task, LukePi (without Node degree classification task) and LukePi (without Edge recovery task drops slightly behind the LukePi (without pre-training) in weak cold start scenario, but achieves on average 2.85% and 3.75% performance boosts in cold start scenario. This result suggests that pre-trained models, which are based on different self-supervised learning objectives, could have a bias to capture specific data patterns, thus lacking the ability to extracts features comprehensively. However, the full LukePi achieves the best performance in all scenarios, which confirms that integrating the two tasks leverages the strengths of each task can learn a more comprehensive and effective embeddings, making the model to achieve good performance across scenarios.

**TABLE V.** Performance of LukePi and three variants on SL prediction task.

| Model | Weak cold start (AUPR) | Cold start (AUPR) |
| --- | --- | --- |
| w/o pre-training | 77.04±1.46 | 69.75±0.04 |
| w/o node degree classification task | 75.37±0.01 | 72.60±6.04 |
| w/o edge recovery task | 74.93±0.67 | 73.50±6.18 |
| LukePi | **78.89±1.04** | **74.36±4.85** |

**TABLE VI.** Performance of LukePi and three variants on DTI prediction task.

| Model | Weak cold start (AUPR) | Cold start (AUPR) |
| --- | --- | --- |
| w/o pre-training | 65.77±3.93 | 70.62±3.50 |
| w/o node degree classification task | 70.60±4.07 | 64.70±8.86 |
| w/o edge recovery task | 70.86±3.86 | 69.52±5.70 |
| LukePi | **77.62±1.98** | **78.22±6.59** |

### E. Case study

To further assess the potential of LukePi in predicting real-world biological pairwise interactions, we apply LukePi to predict SL gene pairs in ovarian cancer (OC). LukePi is fine-tuned using the SynLethDB dataset and then tested on an independent SL dataset obtained through high-throughput screening [9], where these gene pairs are specifically excluded from the training data to prevent data leakage. Additionally, we use the experimental scores [9] to select the most confident negative samples. The final OC test dataset includes 15 SL pairs and 60 negative (non-SL) pairs. Next, we analyze the predictions of LukePi on this OC test data.

#### 1) Prediction of SL gene pairs in ovarian cancer

First, LukePi achieves an overall AUC value of 0.74, indicating a good concordance between the predicted and true SL labels. Secondly, LukePi assigns higher SL prediction scores to positive SL gene pairs (median 2.19) compared to non-SL pairs (median 0.26) with a Wilcoxon test p-value of 5.7 × 10^−3^, as shown in Figure 6A. We rank all the gene pairs based on their prediction scores generated by LukePi, as illustrated in Figure 6B. We then investigate the top 15 candidate gene pairs, with the literature evidence presented in Supplementary Table S3. LukePi successfully predicts 5 SL pairs among these top candidate gene pairs. Notably, we find that LukePi identifies two well-known SL gene pairs in ovarian cancer: BRCA1-PARP1 (ranked 1st) and ARID1A-EZH2 (ranked 5th). These results suggest that LukePi provides an effective means of ranking gene pairs according to their likelihood of being SL.

**Fig. 6.**
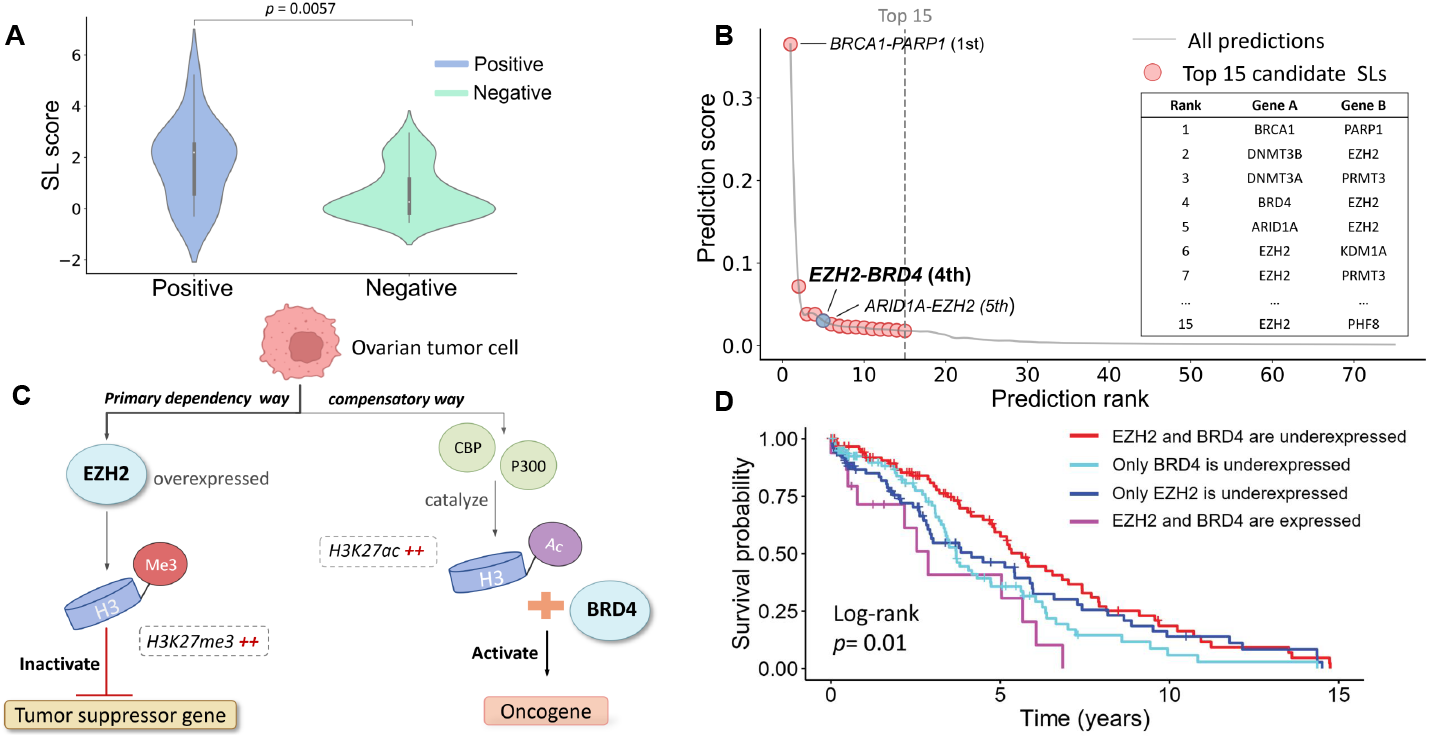
Analysis of SLs predicted by LukePi from an ovarian cancer dataset. (A) Distribution of SL scores for positive versus negative SL pairs, with the p-value calculated using a Wilcoxon rank sum test. (B) Prediction scores versus rank for gene pairs on the SL ovarian cancer dataset. The top 15 predictions (red dots) are located to the left of the dotted gray line, with BRCA1-PARP1 and ARID1A-EZH2 previously validated as SL pairs. EZH2-BRD4 is identified as promising SL pair in ovarian cancer. (C) Overview of the potential biological mechanism underlying the synthetic lethality between EZH2 and BRD4 in ovarian cancer. (D) Kaplan-Meier curves and Log-rank test p-values showing survival differences based on EZH2 and BRD4 expression levels in ovarian cancer patient cohorts.

#### 2) EZH2-BRD4 is a promising SL pair in ovarian cancer

Following the validation of LukePi’s performance in predicting cancer-specific SL pairs, we conduct a literature survey and find that EZH2-BRD4 (ranked 4th) is a promising SL pair, potentially overcoming resistance caused by inhibiting EZH2 alone in ovarian cancer. EZH2, a crucial subunit of the PRC2 complex, is frequently overexpressed in ovarian cancer and regulates the trimethylation of H3K27 (H3K27me3) [3].

However, inhibiting EZH2 alone does not eliminate tumor cells as expected [16], [33]. One possible explanation is that inhibiting EZH2 reduces H3K27me3 but simultaneously triggers a compensatory increase in H3K27ac, as illustrated in Figure 6C. BRD4, a member of the BET protein family, specifically interacts with H3K27ac via its bromodomains, and this interaction is thought to facilitate the transcriptional activation of several oncogenes, thereby promoting the activation of tumor-related pathways, such as the Wnt signaling pathway. This mechanism allows tumor cells to circumvent the therapeutic effects of EZH2is [16]. Consequently, co-inhibiting EZH2 and BRD4 may induce synthetic lethality in ovarian cancer by targeting both primary and compensatory pathways. Moreover, the combination of EZH2 and BRD4 inhibitors has shown tolerable toxicity [16].

To further evaluate the clinical relevance of BRD4 and EZH2, we analyze gene expression and 15-year survival data from 301 ovarian cancer patients in the TCGA database [35]. We hypothesize that the co-underexpression of EZH2 and BRD4 increases tumor vulnerability and improves prognosis. To test this, we classify patients into four groups based on gene expression: those with co-underexpression of both genes and those expressing at least one of these genes. We then plot Kaplan-Meier (KM) curves for each group. Figure 6D shows the survival analysis. Notably, the KM curve for patients with co-underexpression of BRD4 and EZH2 (red line) is significantly higher than that of other groups, with a log-rank p-value of 0.01, indicating longer survival times. These findings suggest that EZH2 and BRD4 could be a promising therapeutic pair in ovarian cancer, though further experimental validation is needed.

Generally, this case study indicates that LukePi can effectively identify well-known SL pairs and uncover promising SL pairs, demonstrating its potential in discovering real-world biological interactions.

## V. Discussion and Conclusion

Pairwise interaction prediction plays a crucial role in understanding various biological processes. However, it remains challenging due to the lack of sufficient labeled data. This limitation particularly affects the performance of supervised GNN-based models. In this paper, we propose LukePi, a self-supervised pre-training framework, for predicting diverse biological pairwise interactions. LukePi guides a GNN model in capturing knowledge from a BKG by a novel and expressive combination of two self-supervised tasks: topology-based node degree classification and semantics-based edge recovery. Specifically, the node degree classification requests the GNN model to predict the degree class label of a node based on the topological structure of its neighborhood in the BKG, forcing LukePi to capture the topological context of a node from the graph. Meanwhile, the edge recovery task enhances the understanding of contextual knowledge by recovering the semantic features of interactions from a randomly disrupted BKG. Leveraging the two self-supervised tasks, LukePi captures shared biomedical knowledge across various types of interactions from a BKG and then produce effective node representations. After pre-training, LukePi can be fine-tuned for making task-specific predictions. To assess the performance of LukePi, we conducted extensive experiments on two important pairwise interactions, i.e. SLs and DTIs. Evaluated in the distribution-shift (i.e., weak cold start and cold start scenarios) and low-data scenarios, LukePi consistently outperforms several state-of-the-art methods, demonstrating its superior performance. Furthermore, the SL prediction by LukePi could uncover novel SL-based anti-cancer drug targets. Our case study suggests that BRD4 and EZH2 predicted by LukePi as a top candidate SL gene pair could be a promising drug target for treating ovarian cancer. In summary, these results demonstrate that pre-training on BKGs is a promising strategy for accurately predicting biological interactions in scenarios with limited labeled data. The source codes and datasets curated in this study are available on GitHub at https://github.com/JieZheng-ShanghaiTech/LukePi.

Although LukePi has demonstrated superior predictive performance, it still has limitations to be addressed in the future. One limitation is that it relies heavily on knowledge graphs, which could limit the model’s ability to generate comprehensive representations of biomolecular entities. In the future, we aim to integrate other modalities of biological data, such as transcriptomics data and protein sequence data, to further enhance the performance of LukePi.

