## Supplemental Material for "Learning universal knowledge graph embedding for predicting biomedical pairwise interactions"

#### 1 Baselines

First, we evaluate five GNN models: GCN [1], GAT [2], GIN [3], RGCN [4], and HGT [5]. GCN generates node representations using symmetric normalized aggregation and self-loop updates. GAT employs an attention mechanism to capture the correlation between a central node and its neighbors. GIN aims to generate node representations that are consistent across isomorphic graphs. RGCN introduces relation-specific weight matrices to model different types of relations in the graph convolutional layer. HGT generates dedicated representations for different types of nodes by designing node- and edge-type dependent parameters for heterogeneous attention. GCN, GAT, and GIN are designed for homogeneous graphs, while RGCN and HGT are tailored for heterogeneous graphs.

Second, we introduce three self-supervised pre-training methods: EdgePred [6], EdgeMask [6], and GCC [7]. EdgePred randomly samples pseudo and true edges from a graph as negative/positive samples, and lets the GNN classify whether candidate edges are true in the graph. EdgeMask masks node/edge features and lets the GNN predict these attributes. GCC adopts sub-graph instance discrimination as a pre-training task.

Third, we evaluate seven task-specific methods: KG4SL [8] and PT-GNN [9] for the SL task, ELISL [10] and MVGCN-iSL [11] for the LAML-specific SL task; and GraphDTA [12], DrugBAN [13], and MSSL2drug [14] for the DTI task. Specifically, KG4SL uses a GCN to learn gene information from a KG and predict SL interactions. PT-GNN leverages rich biological data, including PPI and gene ontology data, to pre-train GCN models and then transfer to SL prediction. MVGCN-iSL is a GNN-based model that predicts SL interactions by integrating five biological networks to learn gene embeddings specific to a cell line or cancer type. ELISL uses multiple features such as gene expression data, mutation data, and clinical data to characterize gene pairs, employing a random forest model to predict SLs for individual cancers. For DTI prediction, GraphDTA proposes a GNN-based method for drug-target binding affinity prediction. DrugBAN captures pairwise local interactions between drugs and targets using a bilinear attention mechanism to predict drug-target interactions. MSSL2drug proposes multitask joint strategies of self-supervised learning on biomedical networks for drug discovery [14].

#### 2 Experimental datasets

In this section, we introduce the details about the pre-training dataset PrimeKG [15]. It integrates 20 high-quality resources to represent ten major biological scales, including disease-associated protein perturbations, biological processes and pathways, anatomical and phenotypic scales, and the entire range of approved drugs with their therapeutic actions, significantly expanding previous efforts in disease-rooted knowledge graphs. It consists of 129,075 nodes and 8,100,498 edges, including 10 types of nodes and 30 types of edges. Table S1 and Table S2 provide the statistical information about nodes and edges in the PrimeKG, respectively.

Table S1: Statistic data of nodes in the PrimeKG.

| **Node Type** | **Count** | **Percent (%)** |
|---|---|---|
| Biological process | 28,642 | 22.1 |
| Protein | 27,671 | 21.4 |
| Disease | 17,080 | 13.2 |
| Phenotype | 15,311 | 11.8 |
| Anatomy | 14,035 | 10.8 |
| Molecular function | 11,169 | 8.6 |
| Drug | 7,795 | 6.2 |
| Cellular component | 4,176 | 3.2 |
| Pathway | 2,516 | 1.9 |
| Exposure | 818 | 0.6 |
| Total | 129,375 | 100.0 |

Table S2: Statistic data of edges in the PrimeKG.

| **Relation** | **Percent(%)** |
|---|---|
| Anatomy - Protein (present) | 37.5 |
| Drug - Drug | 33 |
| Protein - Protein | 7.9 |
| Disease - Phenotype (positive) | 3.7 |
| Biological process - Protein | 3.6 |
| Cellular component - Protein | 2.1 |
| Disease - Protein | 2 |
| Molecular function - Protein | 1.7 |
| Drug - Phenotype | 1.6 |
| Biological process - Biological process | 1.3 |
| Pathway - Protein | 1.1 |
| Disease - Disease | 0.8 |
| Drug - Disease (contraindication | 0.8 |
| Drug - Protein | 0.6 |
| Anatomy - Protein (absent) | 0.5 |
| Phenotype - Phenotype | 0.5 |
| Anatomy - Anatomy | 0.3 |
| Molecular function - Molecular function | 0.3 |
| Drug - Disease (indication) | 0.2 |
| Cellular component - Cellular component | 0.1 |
| Phenotype - Protein | 0.1 |
| Drug - Disease (off-label use) | 0.1 |
| Pathway - Pathway | 0.1 |
| Exposure - Disease | 0.1 |
| Exposure - Exposure | 0.1 |
| Exposure - Biological process | <0.1 |
| Exposure - Protein | <0.1 |
| Disease - Phenotype (negative) | <0.1 |
| Exposure - Molecular function | <0.1 |
| Exposure - Cellular component | <0.1 |
| Total | 100 |

Table S3: Performance of LukePi and three variants on SL prediction task. '✓' means the use of the self-supervised task, while '✗' refers to removing the self-supervised task.

| Ablation settings | | Weak cold start | | Cold start | |
|---|---|---|---|---|---|
| Node degree classification | Edge recovery task | AUPR(%) | BACC(%) | AUPR(%) | BACC(%) |
| ✗ | ✗ | 77.04±1.46 | 69.32±1.79 | 69.75±0.04 | 60.44±0.05 |
| ✓ | ✗ | 75.37±0.01 | 64.36±0.06 | 72.60±6.04 | 64.46±6.94 |
| ✗ | ✓ | 74.93±0.67 | 62.49±7.34 | 73.50±6.18 | 65.67±6.87 |
| ✓ | ✓ | **78.89**±**1.04** | **69.46**±**2.52** | **74.36**±**4.85** | **65.99**±**6.27** |

Table S4: Performance of LukePi and three variants on DTI prediction task. '✓' means the use of the self-supervised task, while '✗' refers to removing the self-supervised task.

| Ablation settings | | Weak cold start | | Cold start | |
|---|---|---|---|---|---|
| Node degree classification | Edge recovery task | AUPR(%) | BACC(%) | AUPR(%) | BACC(%) |
| ✗ | ✗ | 65.77±3.93 | 51.80±2.44 | 70.62±3.50 | 55.89±5.45 |
| ✓ | ✗ | 70.60±4.07 | 62.05±4.13 | 64.70±8.86 | 54.58±6.99 |
| ✗ | ✓ | 70.86±3.86 | 61.03±5.10 | 69.52±5.70 | 58.28±4.83 |
| ✓ | ✓ | **77.62**±**1.98** | **67.72**±**3.77** | **78.22**±**6.59** | **68.42**±**6.51** |

### 3 Ablation study

To evaluate the effect of the two self-supervised tasks proposed in LukePi, we conduct an ablation study by creating three model variants: LukePi (without Node degree classification task), which skips the node degree classification task; LukePi (without Edge recovery task), which drops the edge recovery task, and LukePi (without pre-training), which is trained without any pre-training tasks. For both SL and DTI prediction tasks, we assess the performance of these three variants along with the original LukePi under weak cold start and cold start scenarios. We use the same experimental configurations to ensure a fair comparison. The results of the SL task and DTI task are presented in Table S3 and Table S4, respectively. We have the following two main observations based on the results.

First, the performance of two variants drops significantly compared with the full LukePi in each scenario. For example, in the SL prediction task, compared to the full LukePi, the average AUC scores of LukePi (with the node degree classification task) and LukePi (with the edge recovery task) decrease by at least 3.52% and 0.86%, respectively. This decline is even more significant in the DTI prediction task. These results confirm the individual importance of the two pre-training tasks for the performance of LukePi. Secondly, integrating multiple pre-training tasks allows the model to learn more comprehensive representations, thereby enhancing its ability to adapt to different data distributions and scenarios. As shown in Table S3 and Table S4, only leveraging single pre-training tasks outperform LukePi (without pre-training) in some cases, but the improvement is inconsistent in specific scenarios. For example, for SL task, LukePi (without Node degree classification task) and LukePi (without Edge recovery task drops slightly behind the LukePi (without pre-training) in weak cold start scenario, but achieves on average 2.85% and 3.75% performance boosts in cold start scenario. This result suggests that pre-trained models, which are based on different self-supervised learning objectives, could have a bias to capture specific data patterns, thus lacking the ability to extracts features comprehensively. However, the full LukePi achieves the best performance in all scenarios, which confirms that integrating the two tasks leverages the strengths of each task can learn a more comprehensive and effective embeddings, making the model to achieve good performance across scenarios.

Table S5: The top 15 candidate gene pairs predicted by LukePi for ovarian cancer and their supporting evidence from low-throughput studies.

| Rank | Candidate gene A | Candidate gene B | Evidence |
|---|---|---|---|
| 1 | BRCA1 | PARP1 | [17, 18] |
| 2 | DNMT3B | EZH2 | Unknown |
| 3 | DNMT3A | PRMT3 | Unknown |
| 4 | BRD4 | EZH2 | [19] |
| 5 | ARID1A | EZH2 | [20, 21] |
| 6 | EZH2 | KDM1A | [22] |
| 7 | EZH2 | PRMT3 | Unknown |
| 8 | DNMT3B | ING3 | Unknown |
| 9 | EZH2 | KDM5B | [16] |
| 10 | CREBBP | ING2 | Unknown |
| 11 | DNMT3B | PHF2 | Unknown |
| 12 | EZH2 | KDM5C | Unknown |
| 13 | CREBBP | KDM5A | Unknown |
| 14 | CREBBP | ING3 | Unknown |
| 15 | EZH2 | PHF8 | Unknown |

### 4 Case study

To further validate these results predicted by LukePi, we conduct a literature survey for each candidate gene pair and provide the corresponding low-throughput studies. Table S5 demonstrates the top 15 candidate SL pairs in ovarian cancer. Some candidate gene pairs still lack known supporting evidence. However, this is intuitive, as validated SL gene pairs represent only a small fraction of the total candidate pairs so far. In addition, although evidence for the SL relationship between EZH2 and KDM5B is primarily found in leukemia [16], this validated SL relationship in leukemia encourages us to explore its potential synthetic lethality in ovarian cancer and warrants further investigation to confirm its potential in this context.

[3] Keyulu Xu, Weihua Hu, Jure Leskovec, and Stefanie Jegelka. How powerful are graph neural networks? *arXiv preprint arXiv:1810.00826*, 2018.

[4] Michael Schlichtkrull, Thomas N Kipf, Peter Bloem, Rianne Van Den Berg, Ivan Titov, and Max Welling. Modeling relational data with graph convolutional networks. In *The Semantic Web: 15th International Conference, ESWC 2018, Heraklion, Crete, Greece, June 3–7, 2018, Proceedings 15*, pages 593–607. Springer, 2018.

[5] Ziniu Hu, Yuxiao Dong, Kuansan Wang, and Yizhou Sun. Heterogeneous graph transformer. In *Proceedings of the web conference 2020*, pages 2704–2710, 2020.

[6] W Hu, B Liu, J Gomes, M Zitnik, P Liang, V Pande, and J Leskovec. Strategies for pre-training graph neural networks. In *International Conference on Learning Representations (ICLR)*, 2020.

[7] Jiezhong Qiu, Qibin Chen, Yuxiao Dong, Jing Zhang, Hongxia Yang, Ming Ding, Kuansan Wang, and Jie Tang. Gcc: Graph contrastive coding for graph neural network pre-training. In *Proceedings of the 26th ACM SIGKDD international conference on knowledge discovery & data mining*, pages 1150–1160, 2020.

[8] Shike Wang, Fan Xu, Yunyang Li, Jie Wang, Ke Zhang, Yong Liu, Min Wu, and Jie Zheng. KG4SL: knowledge graph neural network for synthetic lethality prediction in human cancers. *Bioinformatics*, 37(Supplement_1):i418–i425, 2021.
